# Relative distances between homology groups to assess persistent defects in time series

**DOI:** 10.1101/595785

**Authors:** Juan G. Diaz Ochoa

**Affiliations:** PerMediQ GmbH, Pelargusstr. 2, 70180 Stuttgart

**Keywords:** Time Series, Persistent Homology, Machine Learning, Medicine and Biology

## Abstract

It is common to consider a data-intensive strategy to be an appropriate way to develop systemic analyses in biology and physiology. Therefore, options for data collection, sampling, standardization, visualization, and interpretation determine how causes are identified in time series to build mathematical models. However, there are often biases in the collected data that can affect the validity of the model: while collecting enough large datasets seems to be a good strategy for reducing the bias of the collected data, persistent and dynamical anomalies in the data structure can affect the overall validity of the model. In this work we present a methodology based on the definition of homological groups to evaluate persistent anomalies in the structure of the sampled time series. In this evaluation relevant patterns in the combination of different time series are clustered and grouped to customize the identification of causal relationships between parameters. We test this methodology on data collected from patients using mobile sensors to test the response to physical exercise in real-world conditions and outside the lab. With this methodology we plan to obtain a patient stratification of the time series to customize models in medicine.

## Introduction

High dimensional time series is are ubiquitous in several fields, from economics to biology and medicine. They are relevant because its combination helps to find relationships to infer causal relations between observations, for instance in the identification of the driving and driven systems from experimental time series (Paluš and Vejmelka, 2007). This allows the identification of the coupling between different parameters to derive and train descriptive and predictive models. Previous investigations focused on the definition of causality tests using time series (Paluš and Vejmelka, 2007), for instance using transfer entropy (Mao and Shang, 2017). But the understanding of causalities and the successful implementation of models requires a well-founded analysis of the influence of the sampled data^1^, and in some cases *the inference of causality can be complicated by a bias in estimation from a limited amount of possibly noisy data* (Paluš and Vejmelka, 2007).

This inference of causality is based on the notion of cooperative behavior of coupled complex systems, where synchronization and related phenomena has been observed in physical as well as biological systems (Pikovsky, 2008). Indeed, recently it has become very useful to deal biological systems from the point of view of theoretical informatics. For example, a bird flying in a forest *compute* their trajectory *Γ* according to the tree distribution in the environment. Honeybees are also computing and creating a model of their environment (Roper et al., 2017). But also, a cancer cell adapting its response also represented by *Γ* according to changes in its microenvironment, i.e. depending on the acidity of the tissue or even to toxic chemical substances for the cell (Rossetti et al., 2012). Such representations are useful considering that an environment is non-static: a storm can change the distribution of trees in a forest, or the change of a diet can induce substances able to modify microenvironments in tissues affected by cancer cells (Diaz Ochoa, 2018).

We estimate persistent homology groups to qualitatively assess persistent incoherencies and disbalances in the sampled data associated to the trajectories *Γ*. Since this methodology is robust against noise (Emrani et al., 2014), it is best suited to detect persistent defects in the sampled *Γ*′*s*. Such disbalances are more than errors in the sampling of data and can be identified as persistent and inherent characteristics of the trajectories *Γ*. Such qualitative assessment is not only relevant for the optimization of modelling methods, e.g. avoiding expensive training of models, but also to assure the safety in the use of models by recognizing when a sample of data from a biological system or organism can be represented with a common underlying model, or instead requires a customized mathematical representation, which is for instance helpful to determine if personalization of relevant mathematical models is required for diagnose and therapy in medicine.

To test this kind of stratification based on a persistent bias we use the Mhealth dataset^2^, which contains data from patients wearing IoT^3^ sensors connected to internet devices to measure electro cardiograms (ECGs) and acceleration while they perform physical exercise in normal and non-controlled conditions. In the next section we introduce the required mathematical background. In section three we test this concept using the MHEALTH data set, for patients wearing IoT devices measuring EKGs and acceleration when they perform physical exercise. Finally, we discuss the resent results and perspectives.

## Topological methods for the assessment of bias in the sampled data

### Definition of persistent bias

According to the bias-variance decomposition derivation, the error of a model *f̂*, *Error*(*f̂*), is composed of three terms: a bias that depends on the definitions of the modeler, a variance term, and an unavoidable *irreducible error* term and is given by^4^ 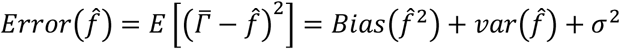,where *Bias*(*f̂* ^2^) is the bias of the model *f̂*.

In this concept the bias is assigned to the way how the data is collected. But the specific reaction to the environment can induce persistent bias in the data structure: the way in which the organism ***k*** autonomously explores its environment, and differs respect other organisms ***l***, has an effect in the variability of the estimated error of a model, such that^5^ (see the appendix 1)

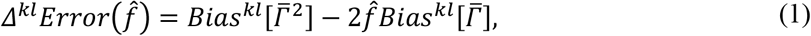

where 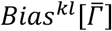 can be defined as (see appendix 1):

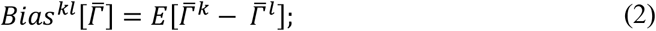

here *E* is the expected value of the difference of the trajectory 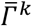 of the organism *k* respect the trajectory 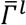 of the organism *l*, 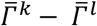. This basically is a perturbation of the error respect the trajectories computed by other organisms generating a response to the environment.

From equation 2 we concentrate us on the analysis of 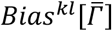, since we want to analyze the relative changes of trajectories between organisms *k* and *l* leading to this variability of the error.

We observe two different cases:

- Considering, the trajectory *Γ*^*k*^ can differ from another sampled trajectory *Γ*^*l*^, the equation 2 represents a persistent bias for each sampled trajectory respect other trajectories and is a representation about how the organism *k* sees the environment, and how this perspective differs from other organisms with their own representation, such that,

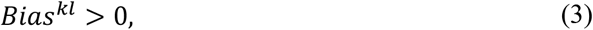

i.e. the organism generates incoherent trajectories respect other organisms.
- On the other hand, when all the trajectories of all the organisms are coherent (for instance the trajectory of a birth in a tree labyrinth is similar to the trajectory of other birds), then

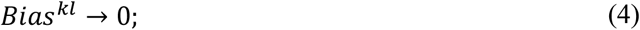

this implies either that the environment is static, or that under high variable environments the organism is rapidly adapting. In the first case the system can be modelled, i.e. there is a function *f̂* representing a trajectory coupled to an environment (which minimizes a given landscape *U*_λ_). This allows the modelling of the trajectory 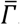, assuming that 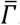 is an observable that somehow represents a reaction to its environment.

Thus, this persistent bias is a natural element in the perturbation of *Error*(*f̂*). This implies that the definition of a valid and stable model does not only depends on the ability of the modeler selecting the best possible model, but also depends on the coherence of the data structure.

### Topological persistence: separation of internal bias from statistical error and modelling

To estimate whether *Bias*^*kl*^ → 0 (see equation 4) we need to assess the structure of the sampled data. The strategy we propose is to assess the topological structure of the data before a model or regression is performed, ideally combining different trajectories in a phase space.

The strategy selected in this work consists in the construction of point clouds 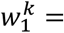 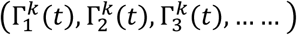 generated from the normalized trajectory sample {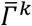} of the organism or system *k*. A point cloud includes a large but finite set of points sampled from a primary form. In the theory t is well known that the combination of the time series of the trajectories 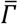, including the time delay of time series, can recover the dynamics of the system (Takens, 1981). Furthermore the presence of harmonic structures in the data, related to this dynamics, can be explored analyzing homological persistence (Emrani et al., 2014). Persistent homology, a tool in algebraic topology, is particularly useful in these situations where the “scale” is not known a priori. It may be viewed as a generalization of hierarchical clustering for higher order topological features, which provides a summary using a bar plot, which essentially validates the different groups discovered using topological invariance (Emrani et al., 2014)^6 7^.

Since *Γ*^λ^ owns a topology that reflects the periodic behavior of a signal with Euler characteristics, this means *Γ*^λ^ owns a function *g* with a compact subset of ℝ^*D*^ and *d*_*k*_: ℝ^*D*^ → ℝ the distance function of *K*. In the following we employ a similar notation as defined by Fasy et. al. (Fasy et al., 2014) for the persistence bars. Consider the *set L*_*t*_ = {*x*: *d*_*u*_(*x*) ≤ *t*} and *k* = *L*_0_. As *t* varies from 0 to ∞, the set *L*_*t*_ changes. *L*_*t*_, defined as a persistent bar, is characterized with persistent topology and summarizes how the topological characteristics *L*_*t*_ change as a function of *t*. Key topological features include zero and first order topology (see figure 1).

**Figure 1.**
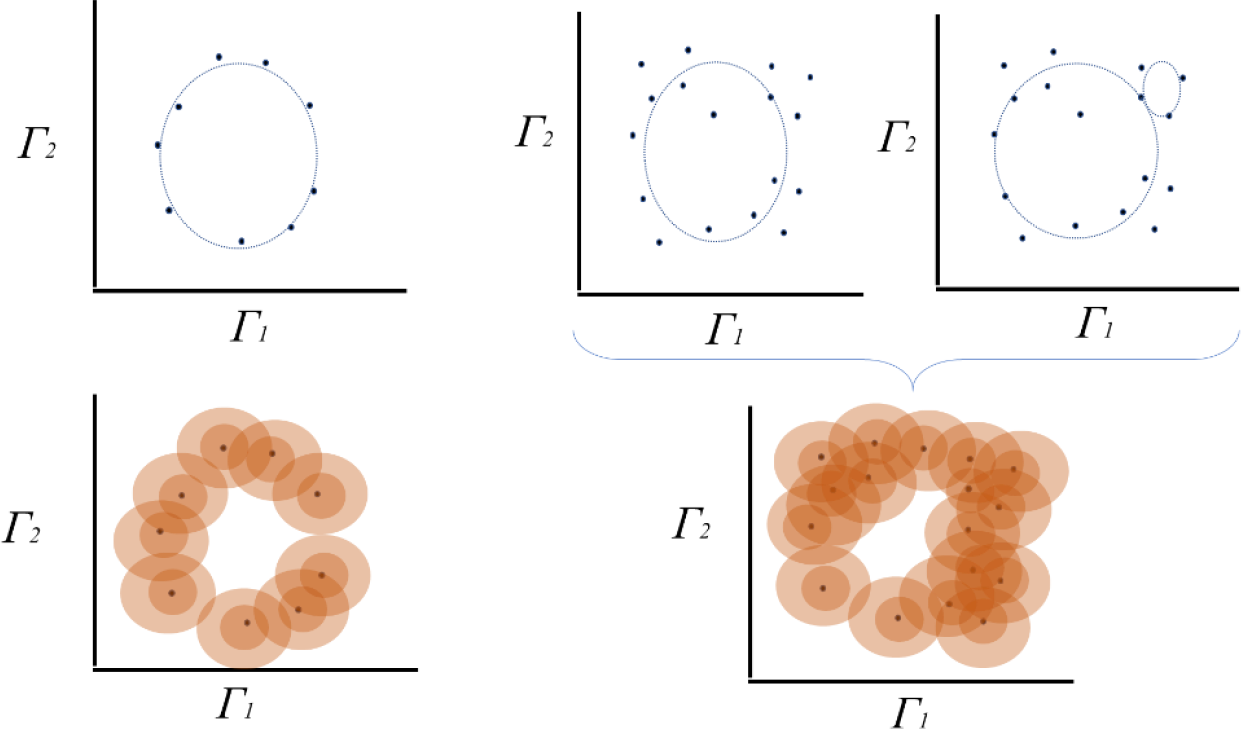
Estimation of topological persistence. In the example left connected points and a surface with homology 0 can be extracted. In the right the same procedure allows the discovery of a second structure with homology 1.

**Figure 2.**
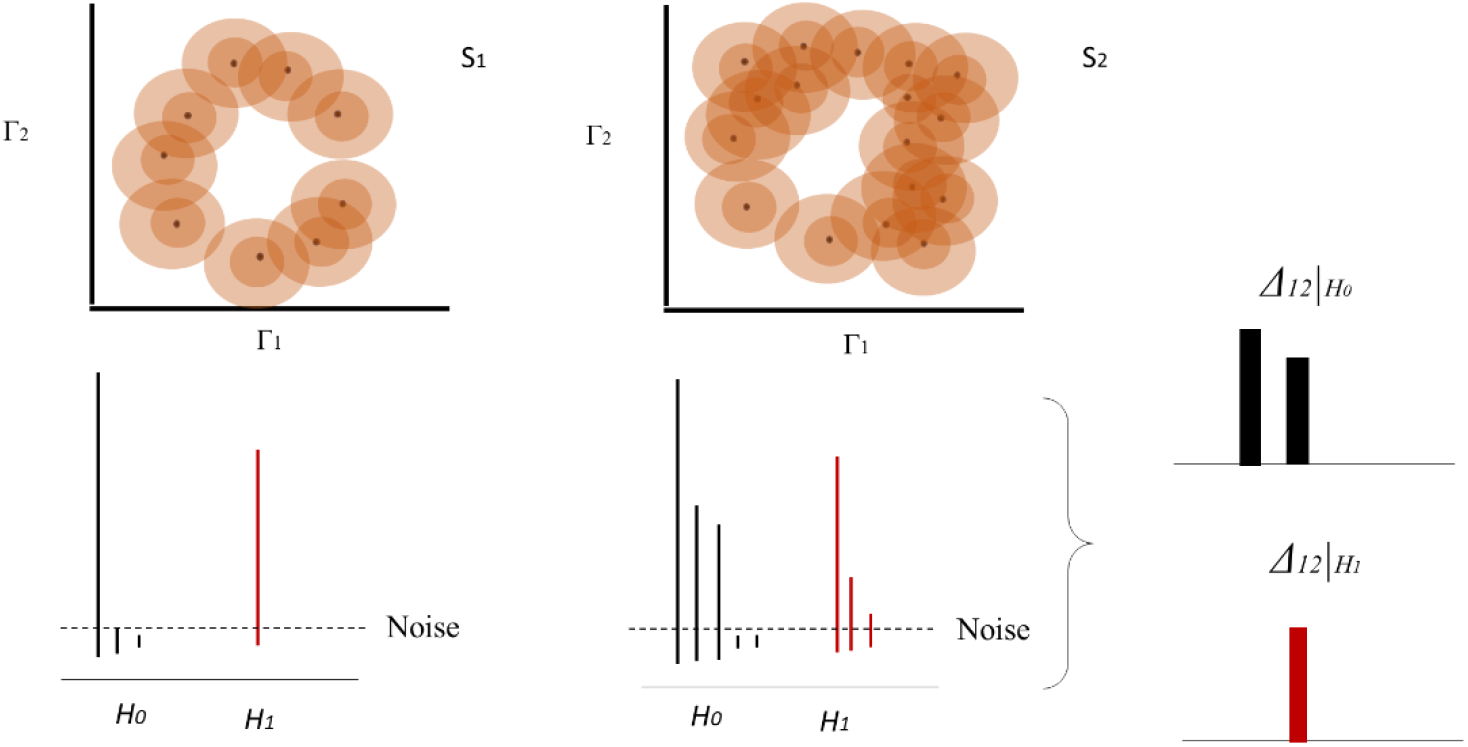
Homology group and bar diagram for two different data sets based on example in figure 1. For S1 the persistent dyagram delivers a homology group H_1_, i.e. a group with genus =1. For S2 the increase in the number of points allows the defintion of a homology gropup H_0_, i.e. a surface with connected points. On the left we schematically represent the difference between both persistent diagrams (for each homology group)

For λ = 1,2, ⋯ *⋀*, if there is a sample of trajectories generating a point cloud {*W*^λ^} then L_t_{w^λ^} are different topological persistent characteristics of the trajectory *Γ*^λ^, such that when equation 3 is combined then,

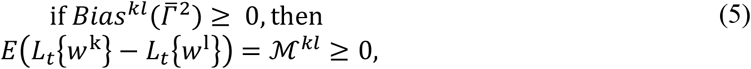

which is equivalent to low change of *Γ*^*k*^ respect *λ* implying a preservation of the internal structure 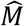 used to compute the trajectory (persistent internal bias). Otherwise,

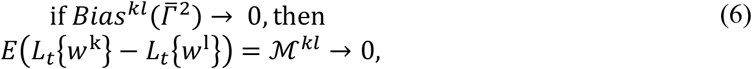

i.e. organisms show similar patterns or a similar response, i.e. 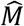 is equivalent for each organism. The matrix 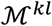 will be called in what follows distortion matrix.

The goal is to make a relative assessment of the topological persistence of a cloud of points. Persistent characteristics as well as pure noise are not of central interest. Instead, persistent topological characteristics, computed as the difference of the persistent bars, or equivalently as the difference between homology groups, provide information about similarity between organisms as well as potential internal bias. <u>Accordingly</u>:

> *low relative persistence of data, i.e. 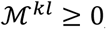, implies a high probability that a customized model f̂_*k*_ is required, i.e. that f̂_*l*_ will probably not completely fit the sampled data of k*

In figure 3 we describe the main steps of this algorithm: each individual generates her own set of trajectories that produce a point cloud *W*^λ^. Using topological homology analysis, the homology is computed for each group, and the persistence is computed using a cluster analysis for each point cloud. This computation essentially extracts the fingerprint from the data generated from each individual. Finally, the relative distance between these fingerprints, i.e. distance between homology groups, 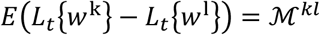 is estimated. Due that we compute a difference of the silhouette of the clustering analysis, the final value 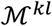 is better represented by a histogram. For this reason, a box plot is used to represent the distribution of 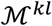 (see appendix 2). The skewness of this distribution provides relevant information about the stability of the perturbation at 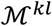.

**Figure 3.**
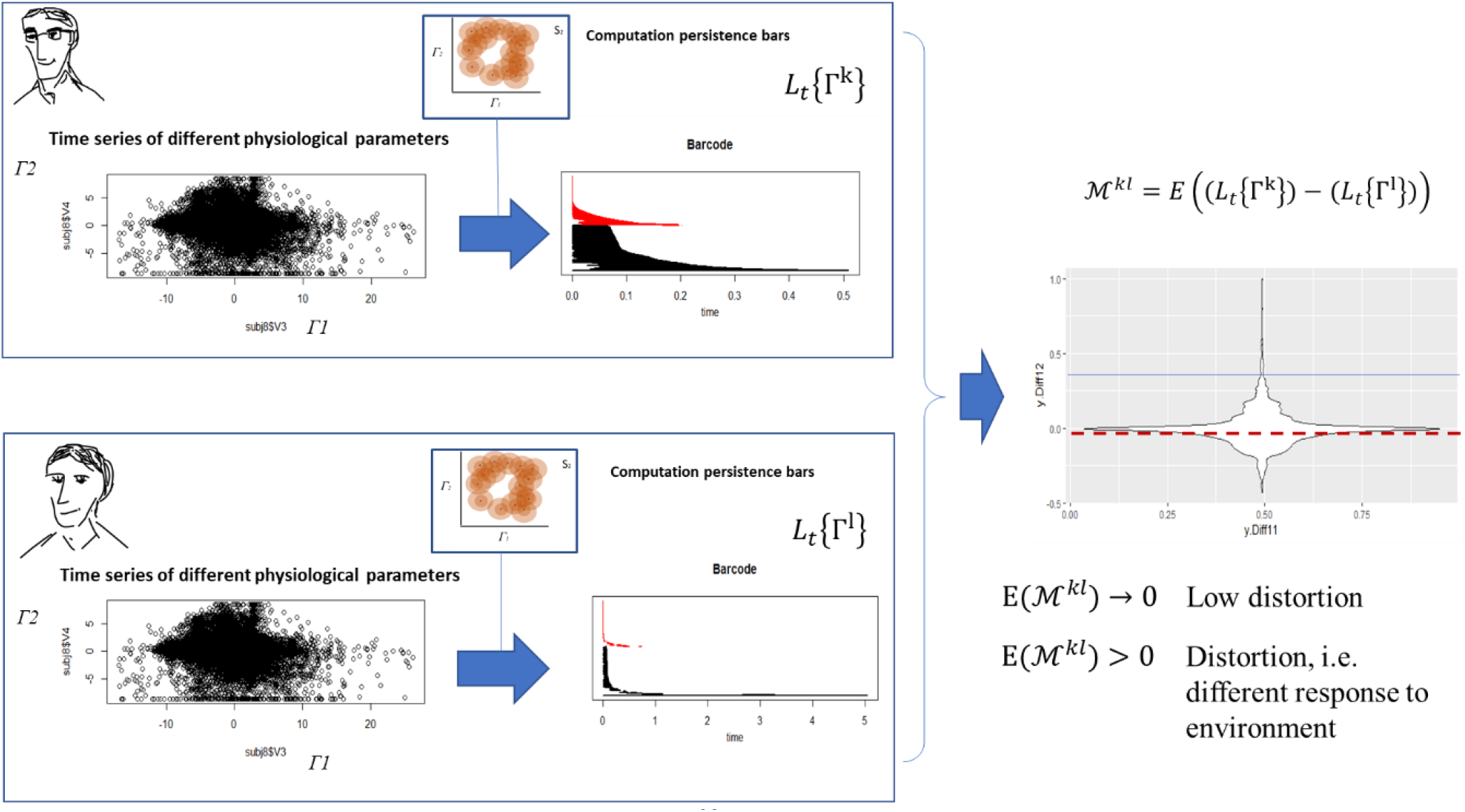
Different steps in the analysis of the matrix 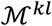 according to equation 8. For example, the data from two individuals is analyzed as a point cloud in a phase space using topological invariance. Once the persistent diagrams are obtained they are normalized respect the highest value of the persistence diagram. The final normalized distribution of the persistence diagram of each individual is subtracted. If the expected value is about 0, then the distortion is minimal, i.e. there is no significant deviation between the patterns of two individuals. Otherwise, a distortion can be identified. Since we deal with the difference between distributions, we employ box plots or violin diagrams to estimate the relative difference between two individual patterns.

## Analysis of patient data generated from IoT

We test the current method in the analysis of physiological signals of patients. As has been suggested in other studies in animals, the locomotor activity is associated to an integrated increase in the activity of different physiological signals (like heart rate, arterial pressure, etc.) (Miyata et al., 2016). Furthermore, the heart response to exercise (macroscopic scale) has origin in complex molecular mechanisms; *for instance in subjects undergoing investigation for angina, individuals with a low chronotropic index (a measure of heart rate response that corrects for exercise capacity) had impaired endothelial function, raised markers of systemic inflammation, and raised concentrations of N‐terminal pro‐brain natriuretic peptide (NT‐proBNP) compared to those with a normal heart rate response* (Routledge and Townend, 2006).

Based on this concept we assume that each individual generates a unique pattern for this integrated response, due to the individual capacity -across several scales- to adapt and/or accommodate to changes in the environment, similar to the case presented in figure 3 (in this case the response to physical exercise).

The data used in this analysis has been obtained from the MHEALTH (Mobile HEALTH) dataset, which comprises body motion and vital signs recordings for ten volunteers of diverse profile while performing several physical activities. Sensors placed on the subject’s chest, right wrist and left ankle are used to measure the motion experienced by diverse body parts, namely, acceleration, rate of turn and magnetic field orientation. The sensor positioned on the chest also provides 2-lead ECG measurements, which can be potentially used for basic heart monitoring, checking for various arrhythmias or looking at the effects of exercise on the ECG. The activities were collected in an out-of-lab environment with no constraints on the way these must be executed, with the exception that the subject should try their best when executing them^8^ (see figure 4). The final raw data is analyzed, i.e. features are extracted from the time series and the main qualitative features are analyzed using a heat map (see appendix 2 for an explanation about how the heat maps in figure 5 are constructed).

**Figure 4.**
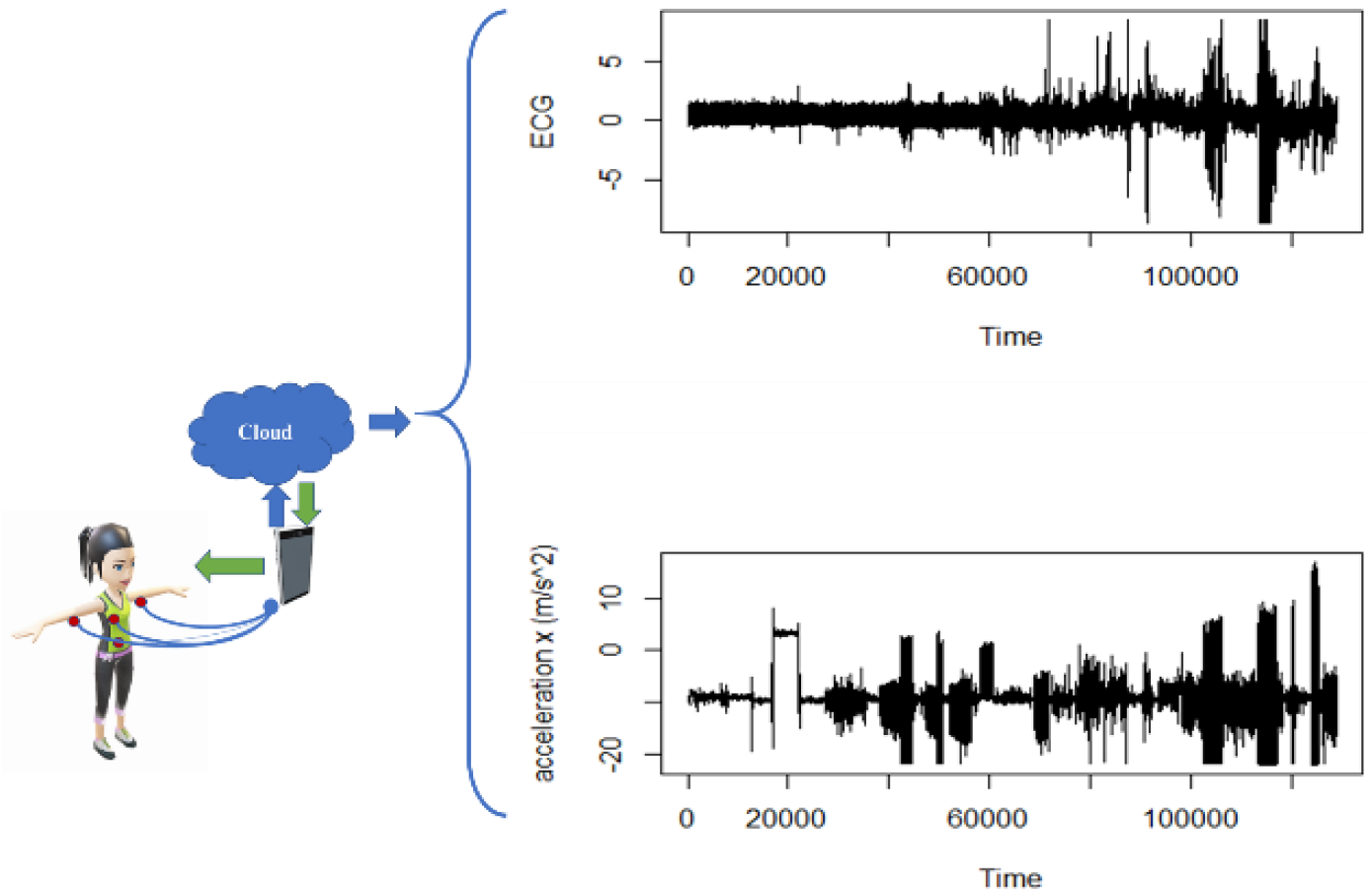
Exemplary data obtained from patients wearing sensors to measure ECG and acceleration in three axes (x, y, z)

**Figure 5.**
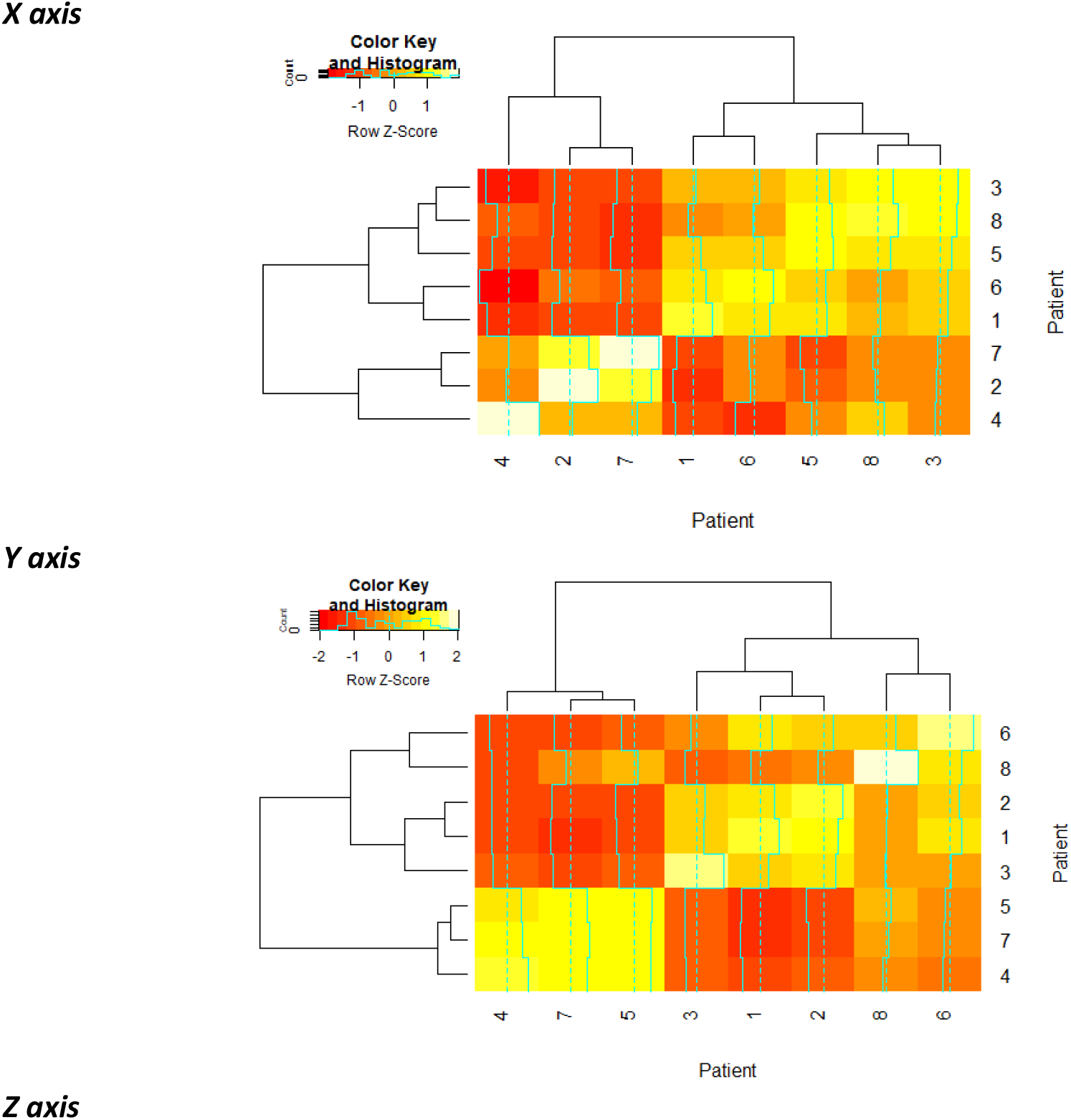
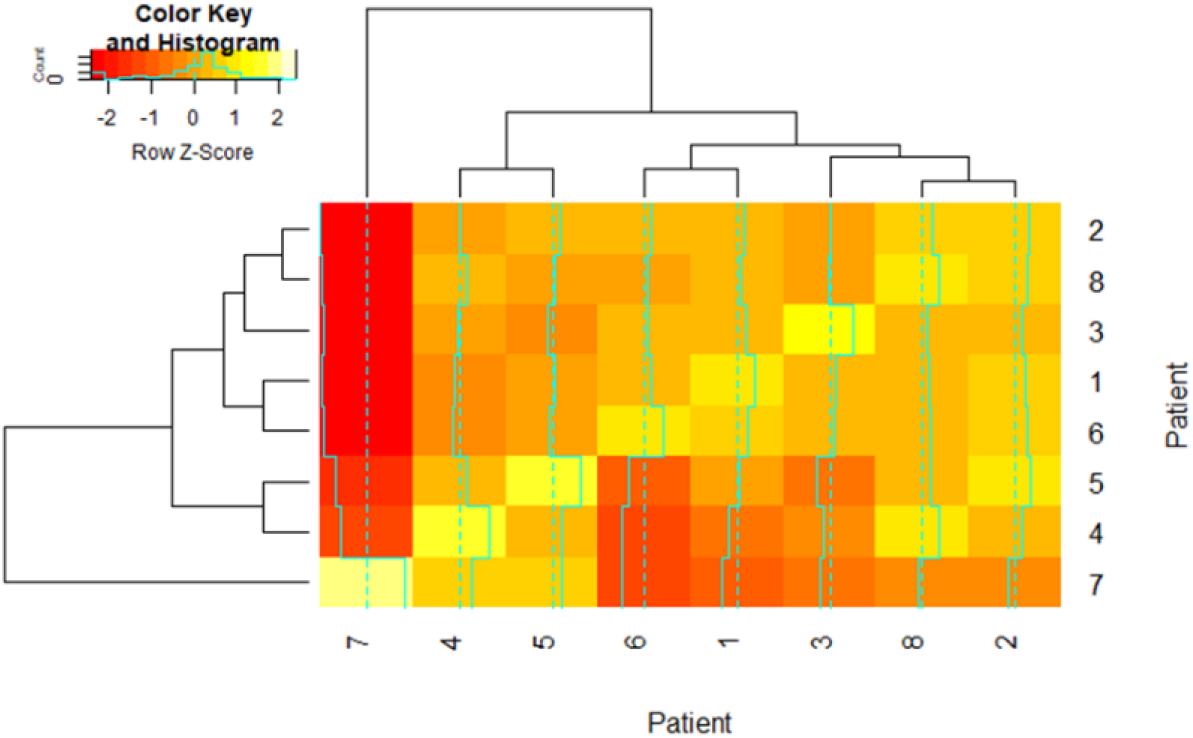
Application of the methodology introduced in fig. 3 for the analysis of physiological data of different patients: once the ECG and acceleration data of a patient doing physical exercise is acquired by IoT devices and stored for instance in the cloud. The correlation of the relative difference of the homology groups (appendix 2) is analyzed for four different patients considering that the acceleration is measured on the x axis (B.), y axis (C.) and z axis (D).

The relative distance between the homology groups 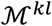 is shown in detail in appendix 2. To facilitate the analysis of these results, the correlation between values have been estimated using a Pearson product-moment correlation coefficient (figure 5).

From this analysis we perform an analysis linking the acceleration measured with a chest sensor in relation to the ECG, and apply the methodology described in the previous section and in figure 3. We observe some similarities, while a rich structure is revealed when the analysis is extended to each axis where the acceleration is measured. A relevant result is that the structure of the clustering is almost equivalent for the three axes, implying that the method delivers a consistent stratification of the individuals.

With the dendrograms analysis we observe the formation of clusters in the homology groups for the patients 7 and 4, implying that there is a correlation between both probands, and in particular that in both cases the relative distance between their homology groups is low. Otherwise 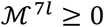 as well as 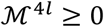 (for the patient 5 this affirmation is non conclusive), or according to equation 5, 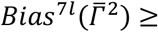 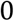 and 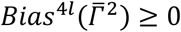. Hence, any inference of causal relations between observations must be differentiated between the patients 7 and 4 and the rest of the individuals in this dataset.

The study is highly qualitative and is reduced to the quantification of relative differences of the data structure in the heatmap, keeping in mind the use of this application for fast data assessments. However, all the assessment delivers more data than the simple qualitative result using homology groups, since it is basically a kind of unsupervised machine learning detecting structures in the time series, which can be used for the computation of prognoses. The use of this methodology will be further explored in next studies.

## Conclusion

We have introduced a qualitative method to assess persistent bias in sampled data. Instead of cumulating and managing very large data sets, it seems to be a better strategy to first recognize which data collections are appropriate and balanced to train models that can be validated and are reliable for further extrapolation. This means, an appropriate customization of models ab initio after assessing persistent bias is more efficient than the training of universal models on several data sets that will be problematic in its validation (Taniguchi et al., 2018)^9^.

To deal with this problem it could be of great advantage to measure the complexity of the data to be analyzed, using for instance a Kolmogorov or a Chaitin complexity measurement (Adriaans, 2018), together with the design of alternative learning architectures. We think that more comprehensive methods and techniques for machine learning are required. The problem is not only the risk that biases and disbalanced information get into big-data sets used to train machines, but it is also the ever-increasing use of resources: the growing processing and storage of information require a lot of energy and resources that end up in form of waste and green-house gases in the atmosphere^10^.

Using concepts of the theoretical informatics and assuming that organisms compute their trajectories after processing information from the environment, and that they adjust to their environment and eventually evolve (Diaz Ochoa, 2018), we develop a method to qualitatively detect this complex behavior by measuring the variability of the modelling error. If the data obtained from any organism’s trajectory has a persistent structure, then the variability of the modelling error is low, implying that a model can be derived and trained. Otherwise, there is not only a bias that can be assigned to the modeler, but also a persistent bias which deteriorates the quality of the statistics.

We suggest that by performing this kind of qualitative analysis before any data processing helps to recognize and introduce convenient data imbalance that not only contributes to a better understanding of the problem (recognition of entities as modelers, and not simply as a blind reaction mechanisms), but also avoids the expensive training of models.

For the detection of persistent structures in trajectories, we have implemented methods using persistent topology for the analysis of time series (Maletić et al., 2016), which have become a promising way to detect patterns in data different to entropy based methods. Despite the implementation of persistent homology is not straightforward, it shows the possibility to better select the data for training and points to the possibility to introduce clever bias in the models to reduce the amount of training data. The concrete application in the analysis of physiological data is a contribution to customize models (for diagnose and therapy) in patients. However, more work is required to standardize and test this methodology in additional datasets.

Thus, we believe that this method is a way of looking at the model through the lens of the training data^11^, and helps to better assess the derivation of causal structures from data for model derivation in medicine and biology, and in general in all the field of Machine Learning.

## Acknowledgments

We are thankful for the critical feedback of Elena Ramirez, as well as useful comments provided by Bernardo Huberman to this manuscript. I also acknowledge the feedback of an anonymous referee.

## Appendix 1

The following is the detailed derivation of the expression linking the variation of error and internal bias:

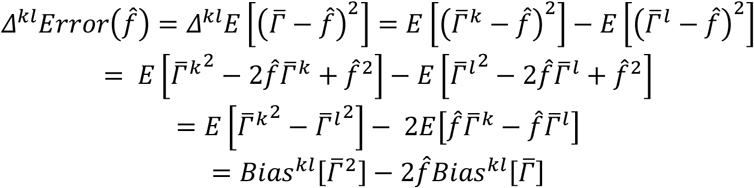

## Appendix 2

Comparation of distortion matrix: in this proposal we employ the following system

- If we compare one patient with himself to assess own changes in the own topological patterns, then we compare the persistence between two different time periods
- If we compare one system against other systems, then we compare the topological persistence of the time series of both individuals in the same time period

In this part we compute the length of the persistence bar as well as the relative difference between the persistence bars. The sampled box plots below (table 1) represents the distribution of the relative differences of persistent features between individuals when accelerations are measured on the *x* axis. Observe that values that deviate from 0 imply that data has an implicit structure that diverge for the specific individual.

**Table A2-1.**
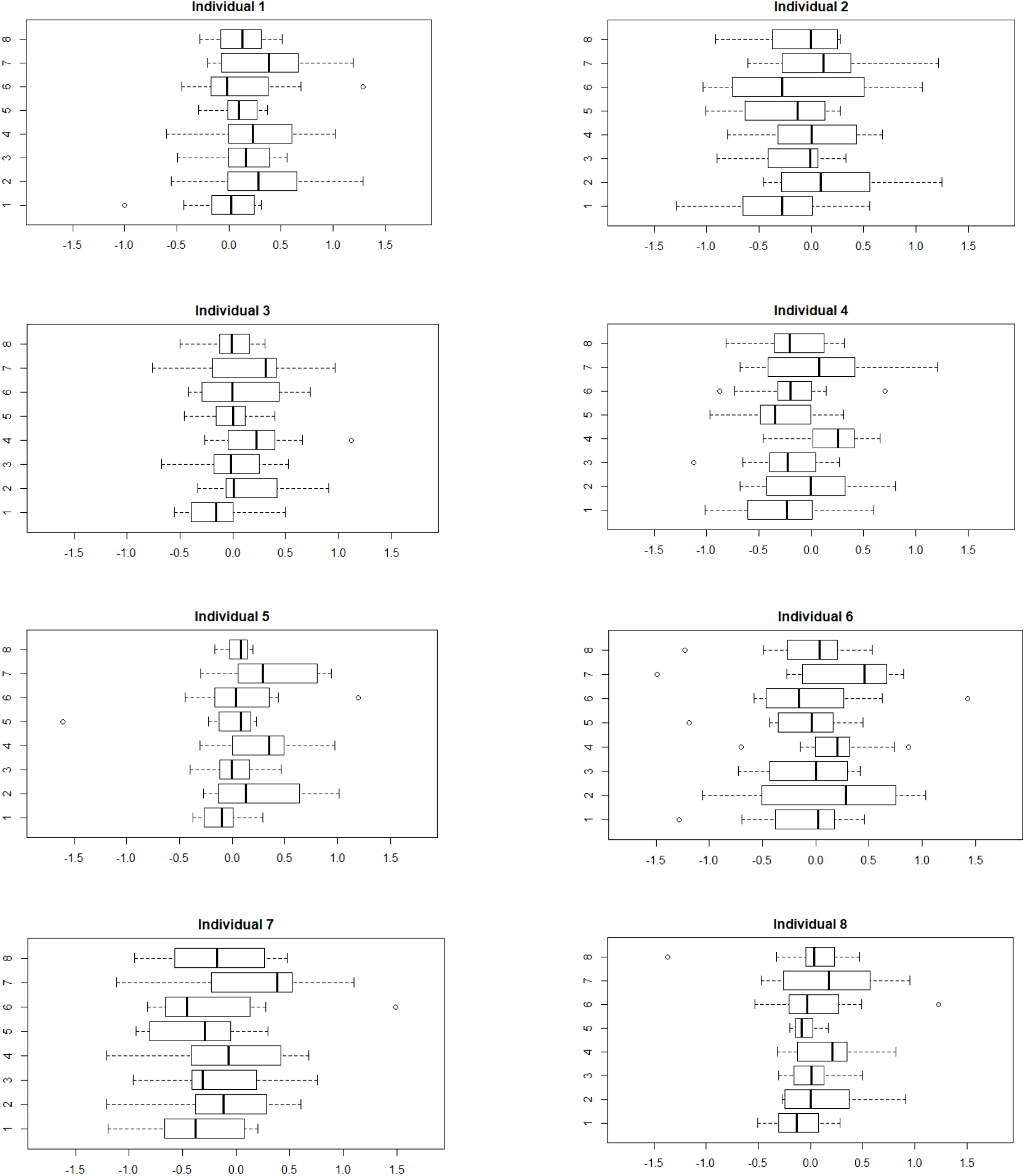
Distribution of the relative distances of the persistence bars between different patients performing physical exercise. Accelerations are measured on the x axis

A comprehensive analysis of this table is made by computing the median of the distributions in the boxplots. We use the median because it is robust against extreme outliers in the distributions presented in the box plots. The final result is presented in the graphic below. Observe that according to this result, individuals 2, 4, and 7 have a different data structure than the rest of individuals, pointing to a persistent bias.

**Figure A2 – 1.**
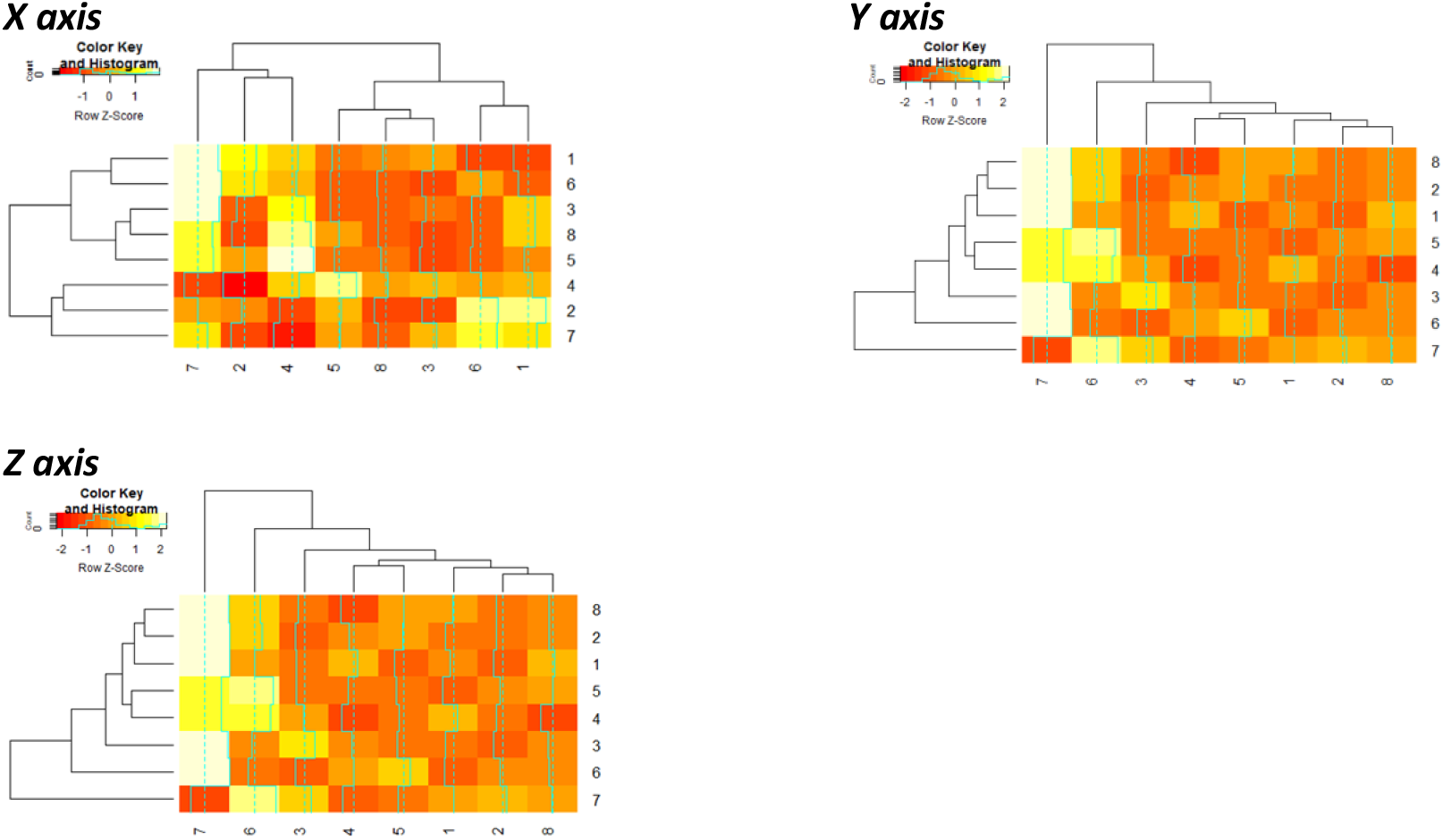
Heatmap for the distribution of the medians of the boxplots from table A2-1

## Footnotes

1 https://arxiv.org/abs/1703.04730v2

2 http://archive.ics.uci.edu/ml/datasets/mhealth+dataset

3 Internet of things

4 http://scott.fortmann-roe.com/docs/BiasVariance.html

6 https://cran.r-project.org/web/packages/TDA/index.html

7 https://cran.r-project.org/web/packages/TDA/vignettes/article.pdf

8 http://archive.ics.uci.edu/ml/datasets/mhealth+dataset Se also this reference for the obtention and use of this data https://www.ncbi.nlm.nih.gov/pmc/articles/PMC4547155/

9 Therefore, we think that any research in machine learning do a better job by dealing with the natural symbiosis between information and life sciences, rather than try to simulate or imitate human cognitive capabilities.

10 Regarding increasing concentration of greenhouse gases in the atmosphere, among them 6% are generated from computation and It, it results extremely important to develop methods to reduce the ecological impact of performing Machine learning.

11 https://arxiv.org/pdf/1703.04730v2.pdf

